# New virus isolates from Italian hydrothermal environments underscore the biogeographic pattern in archaeal virus communities

**DOI:** 10.1101/2020.01.16.907410

**Authors:** Diana P. Baquero, Patrizia Contursi, Monica Piochi, Simonetta Bartolucci, Ying Liu, Virginija Cvirkaite-Krupovic, David Prangishvili, Mart Krupovic

## Abstract

Viruses of hyperthermophilic archaea represent one of the least understood parts of the virosphere, showing little genomic and morphological similarity to viruses of bacteria or eukaryotes. Here, we investigated virus diversity in the active sulfurous fields of the Campi Flegrei volcano in Pozzuoli, Italy. Virus-like particles displaying eight different morphotypes, including lemon-shaped, droplet-shaped and bottle-shaped virions, were observed and five new archaeal viruses proposed to belong to families *Rudiviridae*, *Globuloviridae* and *Tristromaviridae* were isolated and characterized. Two of these viruses infect neutrophilic hyperthermophiles of the genus *Pyrobaculum*, whereas the remaining three have rod-shaped virions typical of the family *Rudiviridae* and infect acidophilic hyperthermophiles belonging to three different genera of the order Sulfolobales, namely, *Saccharolobus*, *Acidianus* and *Metallosphaera*. Notably, *Metallosphaera* rod-shaped virus 1 is the first rudivirus isolated on *Metallosphaera* species. Phylogenomic analysis of the newly isolated and previously sequenced rudiviruses revealed a clear biogeographic pattern, with all Italian rudiviruses forming a monophyletic clade, suggesting geographical structuring of virus communities in extreme geothermal environments. Furthermore, we propose a revised classification of the *Rudiviridae* family, with establishment of five new genera. Collectively, our results further show that high-temperature continental hydrothermal systems harbor a highly diverse virome and shed light on the evolution of archaeal viruses.

## INTRODUCTION

One of the most remarkable features of hyperthermophilic archaea is the diversity and uniqueness of their viruses. Most of these viruses infect members of the phylum Crenarchaeota and are evolutionarily unrelated to viruses infecting bacteria, eukaryotes or even archaea thriving at moderate temperate [1–4]. Thus far, unique to hyperthermophilic archaea are rod-shaped viruses of the families *Rudiviridae* and *Clavaviridae*; filamentous enveloped viruses of the families *Lipothrixviridae* and *Tristromaviridae*; as well as spherical (*Globuloviridae*), ellipsoid (*Ovaliviridae*), droplet-shaped (*Guttaviridae*), coil-shaped (*Spiraviridae*) and bottle-shaped (*Ampullaviridae*) viruses [1,5]. Hyperthermophilic archaea are also infected by two types of spindle-shaped viruses, belonging to the families *Fuselloviridae* and *Bicaudaviridae* [6,7]. Whereas bicaudaviruses appear to be restricted to hyperthermophiles, viruses distantly related to fuselloviruses are also known to infect hyperhalophilic archaea [8], marine hyperthermophilic archaea [9,10] and marine ammonia-oxidizing archaea from the phylum Thaumarchaeota [11]. Finally, hyperthermophilic archaea are also infected by three groups of viruses with icosahedral virions, *Turriviridae* [12], *Portogloboviridae* [13] and two closely related, unclassified viruses infecting *Metallosphaera* species [14]. Portoglobovirus SPV1 is structurally and genomically unrelated to other known viruses [13,15], whereas viruses structurally similar to turriviruses are widespread in all three domains of life [1,16,17]. Structural studies on filamentous and spindle-shaped crenarchaeal viruses have illuminated the molecular details of virion organization and further underscored the lack of relationship to viruses of bacteria and eukaryotes [18–24].

The uniqueness of hyperthermophilic archaeal viruses extends to their genomes, with ∼75% of the genes lacking detectable homologues in sequence databases [25]. All characterized archaeal viruses have DNA genomes, which can be single-stranded (ss) or double-stranded (ds), linear or circular. Comparative genomic and bipartite network analyses have shown that viruses of hyperthermophilic archaea share only few genes with the rest of the virosphere [26]. Furthermore, most of the genes in viruses from different families are family-specific, albeit a handful of genes encoding transcription regulators, glycosyltransferases and anti-CRISPR proteins display broader distribution [27]. These observations, in combination with the structural studies, led to the suggestion that crenarchaeal viruses have originated on multiple independent occasions and constitute a unique part of the virosphere [1].

Although the infection cycles of crenarchaeal viruses has been studied for just a handful of representative viruses, the available data has already provided valuable insight into the virus-host interaction strategies in archaea. For instance, members of the *Fuselloviridae*, *Guttaviridae*, *Turriviridae* and *Bicaudaviridae* are temperate and their genomes can be site-specifically integrated into the host chromosome by means of a virus-encoded integrase [28–33]. In the case of SSV1, the most extensively characterized member of the *Fuselloviridae* family, the virus integrates its genome into the host chromosome at an arginyl-tRNA gene upon cell infection. After exposure to UV light, the excision and replication of the SSV1 viral genomes is induced, leading to virion release [34,35]. Two different egress strategies have been elucidated. The enveloped virions of fusellovirus SSV1 are assembled at the host cell membrane and are released from the cell by a budding mechanism similar to that of some eukaryotic enveloped viruses [36]. By contrast, lytic crenarchaeal viruses belonging to three unrelated families, *Rudiviridae*, *Turriviridae* and *Ovaliviridae*, employ a unique release mechanism based on the formation of pyramidal protrusions on the host cell surface, leading to perforation of the cell envelope and release of intracellularly assembled mature virions [5,37,38].

Single-cell sequencing combined with environmental metagenomics of hydrothermal microbial community from Yellowstone National Park [39] led to the estimation that >60% of cells contain at least one virus type and a majority of these cells contain two or more virus types [40]. However, despite their diversity, distinctiveness and abundance, the number of isolated species of viruses infecting hyperthermophilic archaea remains low compared to the known eukaryotic or bacterial viruses [2]. Indeed, it has been estimated that only about 0.01-0.1% of viruses present in geothermal acidic environments have been isolated [41]. Similarly, using a combination of viral assemblage sequencing and network analysis, it has been estimated that out of 110 identified virus groups, less than 10% represent known archaeal viruses, suggesting that the vast majority of virus clusters represent unknown viruses, likely infecting archaeal hosts [42]. Furthermore, the evolution and structuring of virus communities in terrestrial hydrothermal settings remains poorly understood. Here, to improve understanding on these issues, we explored the diversity of archaeal viruses at the active solfataric field of the Campi Flegrei volcano [43,44] in Pozzuoli, Italy, namely the Pisciarelli hydrothermal area. We report on the isolation and characterization of five new archaeal viruses belonging to three different families and infecting hosts from five different crenarchaeal genera.

## MATERIALS AND METHODS

### Enrichment cultures

Nine samples were collected from hot springs, mud pools and hydrothermally altered terrains at the solfataric field of the Campi Flegrei volcano in Pozzuoli, Italy — the study area known as Pisciarelli — with temperatures ranging from 81 to 96°C and pH between 1 and 7 (Supplementary Information, Table S1). Each sample was inoculated into medium favoring the growth of *Sulfolobus/Saccharolobus* (basal salt solution supplemented with 0.2% tryptone, 0.1% yeast extract, 0.1% sucrose, pH adjusted to 3.5), *Acidianus* (basal salt solution supplemented with 0.2% tryptone, 0.1% yeast extract, sulfur, pH adjusted to 3.5), and *Pyrobaculum* (0.1% yeast extract, 1% tryptone, 0.1-0.3% Na_2_S_2_O_3_ × 5 H_2_O, pH adjusted to 7) species [45,46]. The enrichment cultures were incubated for 10 days at 75°C, except for *Pyrobaculum* cultures, which were grown at 90°C for 15 days without shaking.

### VLP concentration and purification

VLPs were collected after removal of cells from the enrichment culture (7,000 rpm, 20 min, Sorvall 1500 rotor) and concentrated by ultracentrifugation (40,000 rpm, 2h, 10°C, Beckman SW41 rotor). The particles were resuspended in buffer A: 20 mM KH_2_PO_4_, 250 mM NaCl, 2.14 mM MgCl_2_, 0.43 mM Ca(NO_3_)_2_, <0.001% trace elements of *Sulfolobales* medium, pH 6 [47] and visualized under the transmission electron microscope (TEM) FEI Tecnai Biotwin as described below. VLPs were further purified by centrifugation in a CsCl buoyant density gradient (0.45 g ml^−1^) with a Beckman SW41 rotor at 39,000 rpm for 20 h at 10°C. All opalescent bands were collected with a needle and a syringe, dialyzed against buffer A and analyzed by TEM for the presence of VLPs.

### Transmission electron microscopy and VLP analysis

For negative-stain TEM, 5 μl of the samples were applied to carbon‐coated copper grids, negatively stained with 2% uranyl acetate and imaged with the transmission electron microscope FEI Tecnai Biotwin. The width and length of the negatively stained virus particles were determined using ImageJ (n=80) [48].

### Isolation of virus-host pairs

Strains of cells were colony purified by plating dilutions of the *Sulfolobus/Saccharolobus* enrichment cultures onto Phytagel (0.7% [wt/vol]) plates incubated for 7 days at 75 °C. Brownish colonies were toothpicked, inoculated into 1 mL of growth medium and incubated at 75°C for 3 days. To isolate the sensitive hosts of the observed VLPs, 5-μl aliquots of concentrated VLP preparations were spotted onto Phytagel (0.3% [wt/vol]) plates containing lawns of each isolate, as described previously [46]. Inhibition zones were cut out from the phytagel and inoculated into 20 mL of exponentially growing cultures of the corresponding isolates. After incubation at 75°C for 3 days, the VLPs were concentrated by ultracentrifugation (40,000 rpm, 2h, 10°C, Beckman SW41 rotor), resuspended in buffer A and observed by TEM. Pure virus strains were obtained by two rounds of single-plaque purification, as described previously [34].

For *Pyrobaculum* enrichment cultures, a different approach was carried out since it was not possible to obtain single strain isolates. Growing liquid cultures of *P. arsenaticum* 2GA [45], *P. arsenaticum* PZ6 (DSM 13514) [49], *P. calidifontis* VA1 (DSM 21063) [50] and *P. oguniense* TE7 (DSM 13380) [51] were mixed with concentrated VLPs and incubated for 15 days at 90°C. Increase in the number of extracellular viral particles was verified by TEM. In order to obtain cultures producing just one type of viral particles, eight ten-fold serial dilutions (10 ml) of infected cells were established, incubated at 90°C for 22 days and observed by TEM. Because a plaque assay could not be established for VLPs infecting *Pyrobaculum*, we relied on TEM observations for assessment of virus-host interactions.

### Host range

The following laboratory collection strains were used to test the ability to replicate the viruses MRV1, ARV3 and SSRV1: *A. convivator* [31], *A. hospitalis* [52], *Saccharolobus solfataricus* P1 (GenBank accession no. NZ_LT549890) and *S. solfataricus* P2 [53], *Sulfolobus islandicu*s strains REN2H1 [46], HVE10/4 [54], LAL14/1 [55], REY15A [54], ∆C1C2 [56], and *Sulfolobus acidocaldarius* strain DSM639 [57]. CsCl gradient-purified virus particles were added directly to the growing cultures of *Acidianus* species and virus propagation was evaluated by TEM. For *Sulfolobus* and *Saccharolobus* species, spot tests were performed and the replication of the virus was evidenced by the presence of an inhibition zone on the lawn of the tested strains.

The host ranges of PFV2 and PSV2 were examined by adding purified virions to the growing cultures of *P. arsenaticum* PZ6 (DSM 13514) [49], *P. calidifontis* VA1 (DSM 21063) and *P. oguniense* TE7 (DSM 13380) [51]. The virus propagation was monitored by TEM since plaque assays could not be established.

### 16S rDNA sequence determination

DNA was extracted from exponentially growing cell cultures using the Wizard Genomic DNA Purification Kit (Promega). The 16S rRNA genes were amplified by PCR using 519F (5’-CAGCMGCCGCGGTAA) and 1041R (5’-GGCCATGCACCWCCTCTC) primers [58]. The identity of the isolated strains was determined by blastn search of the corresponding 16S rRNA gene sequences against the non-redundant nucleotide sequence database at NCBI.

### Viral DNA extraction, sequencing and analysis of viral genomes

Viral DNA was extracted from purified virus particles as described previously [45]. Sequencing libraries were prepared from 100 ng of DNA with the TruSeq DNA PCR-Free library Prep Kit from Illumina and sequenced on Illumina MiSeq platform with 150-bp paired-end read lengths (Institut Pasteur, France). Raw sequence reads were processed with Trimmomatic v.0.3.6 [59] and assembled using SPAdes 3.11.1 [60] with default parameters. Open reading fragments (ORF) were predicted by GeneMark.hmm [61] and RAST v2.0 [62]. Each predicted ORF was manually inspected for the presence of putative ribosome-binding sites upstream of the start codon. The *in silico-*translated protein sequences were analyzed by BLASTP [63] against the non-redundant protein database at the National Center for Biotechnology Information with an upper threshold E-value of 1e-03. Searches for distant homologs were performed using HHpred [64] against PFAM (Database of Protein Families), PDB (Protein Data Bank) and CDD (Conserved Domains Database) databases. Transmembrane domains were predicted using TMHMM [65]. The sequences of the viral genomes have been deposited in GenBank, with accession numbers listed in Table 2.

### Phylogenomic analysis

All pairwise comparisons of the nucleotide sequences of rudivirus genomes were conducted using the Genome-BLAST Distance Phylogeny (GBDP) method implemented in VICTOR, under settings recommended for prokaryotic viruses [66]. The resulting intergenomic distances derived from pairwise matches (local alignments) were used to infer a balanced minimum evolution tree with branch support via FASTME including SPR postprocessing for D6 formula, which corresponds to the sum of all identities found in high-scoring segment pairs divided by total genome length. Branch support was inferred from 100 pseudo-bootstrap replicates each. The tree was rooted with members of the family *Lipothrixviridae*.

Genome sequences of 19 rudiviruses were further compared using the Gegenees, a tool for fragmented alignment of multiple genomes [67]. The comparison was done in the BLASTN mode, with the settings 200/100. The cutoff threshold for non-conserved material was 40%.

## RESULTS

### Diversity of virus-like particles in enrichments cultures

Nine environmental samples (I1-I9) were collected in hot springs, mud pools and hydrothermally altered terrains of the solfataric fields of the Campi Flegrei volcano in Pozzuoli, Italy, with temperatures ranging from 81 to 96°C and pH values between 1 and 7 (Supplementary Information, Table S1). The enrichment cultures were obtained by inoculating the samples into different media favoring the growth of hyperthermophilic members of the genera *Sulfolobus/Saccharolobus*, *Acidianus* and *Pyrobaculum* [45,46]. Virus-like particles (VLPs) were collected from cell-free culture supernatants and visualized under transmission electron microscope (TEM) as described in Materials and Methods.

A variety of VLPs were detected in the *Sulfolobus/Saccharolobus* enrichment cultures of samples I3 and I9 and the *Pyrobaculum* enrichment culture of sample I4 (Figure 1). Based on virion morphologies, the VLPs detected in the two samples inoculated in *Sulfolobus/Saccharolobus* medium could be assigned to five archaeal virus families: *Fuselloviridae* (Figure 1A), *Bicaudaviridae* (Figure 1B), *Ampullaviridae* (Figure 1C), *Rudiviridae* (Figure 1D) and *Lipothrixviridae* (Figure 1E). VLPs propagated in *Pyrobaculum* medium resembled members of the families *Globuloviridae* (Figure 1F), *Tristromaviridae* (Figure 1G) and *Guttaviridae* (Figure 1H). We next set out to establish pure cultures of these different viruses and to isolate their respective hosts.

**Figure 1.**
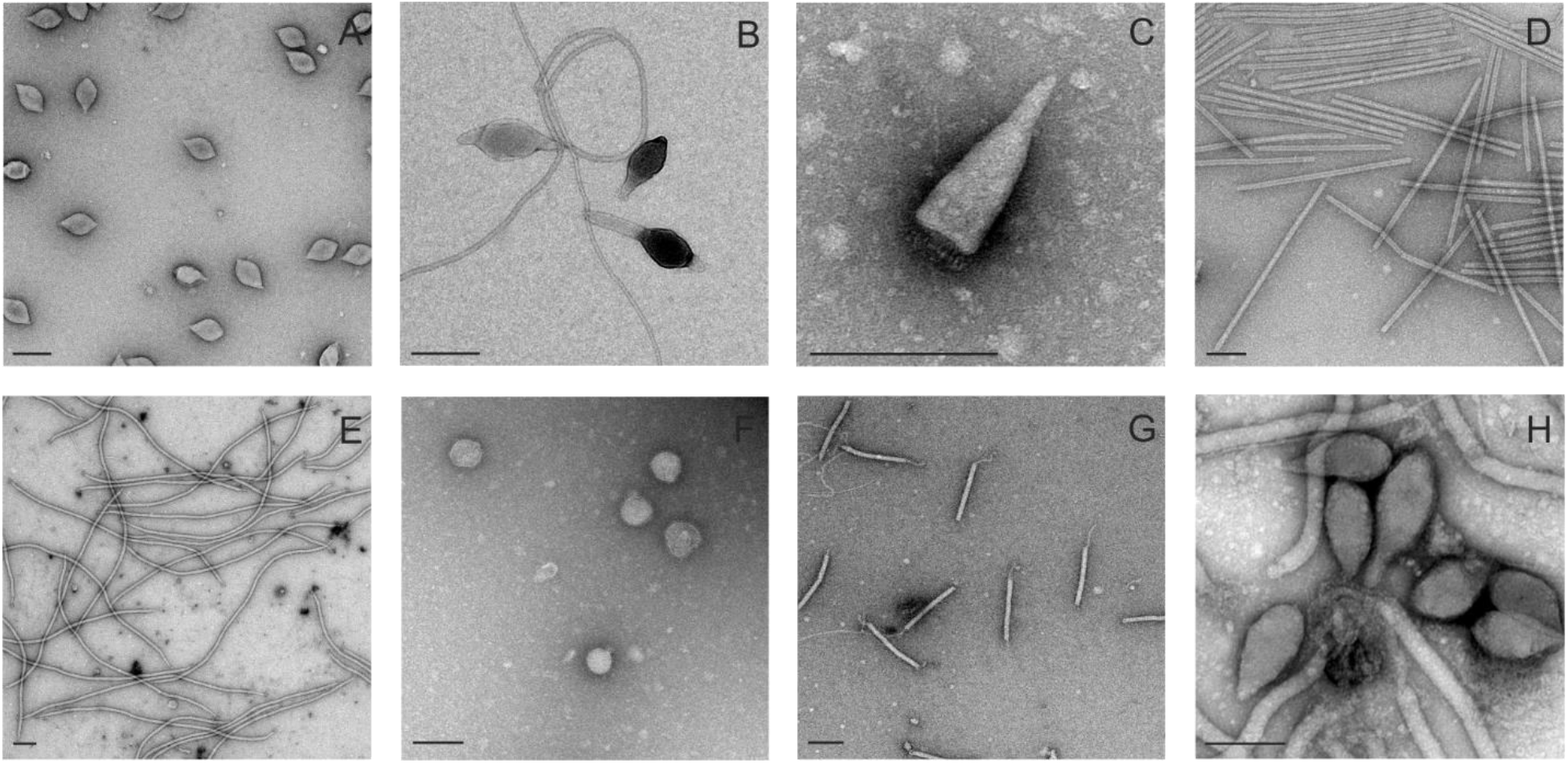
Electron micrographs of the VLPs observed in enrichment cultures: (A) fuselloviruses (tailless lemon-shaped virions), (B) bicaudaviruses (large, tailed lemon-shaped virions), (C) ampullaviruses (bottle-shaped virions), (D) rudiviruses (rod-shaped virions), (E) lipothrixviruses (filamentous virions), (F) globuloviruses (spherical enveloped virions), (G) tristromaviruses (filamentous enveloped virions) and (H) guttaviruses (droplet-shaped virions). Samples were negatively stained with 2% (wt/vol) uranyl acetate. Bars. 200 nm.

### Isolation of virus-host pairs

In order to isolate VLP-propagating strains, 215 single strain isolates were colony purified from the two enrichment cultures established in the *Sulfolobus* medium. Concentrated VLPs were first tested against the isolates by spot test. In case of cell growth inhibition, a liquid culture of the isolate was established and infected with the halo zone observed in the spot test. The production of the viral particles was subsequently verified by TEM. As a result, three strains replicating VLPs were identified. Comparison of their 16S rRNA gene sequences showed that the three strains belong to three different genera of the order Sulfolobales, namely, *Saccharolobus* (until recently known as *Sulfolobus*), *Acidianus* and *Metallosphaera*. The 16S rRNA genes of these strains, POZ9, POZ149 and POZ202, respectively, displayed 100%, 99% and 99% identity to the corresponding genes of *Acidianus brierleyi* DSM 1651 (NZ_CP029289), *Saccharolobus solfataricus* Ron 12/III (X90483) and *Metallosphaera sedula* SARC-M1 (CP012176). Rod-shaped particles of different sizes were propagated successfully in the three isolated strains (Figure 2A-C) and, following the nomenclature used for other rudiviruses, were named *Metallosphaera* rod-shaped virus 1 (MRV1), *Acidianus* rod-shaped virus 3 (ARV3) and *Saccharolobus solfataricus* rod-shaped virus 1 (SSRV1), respectively.

**Figure 2.**
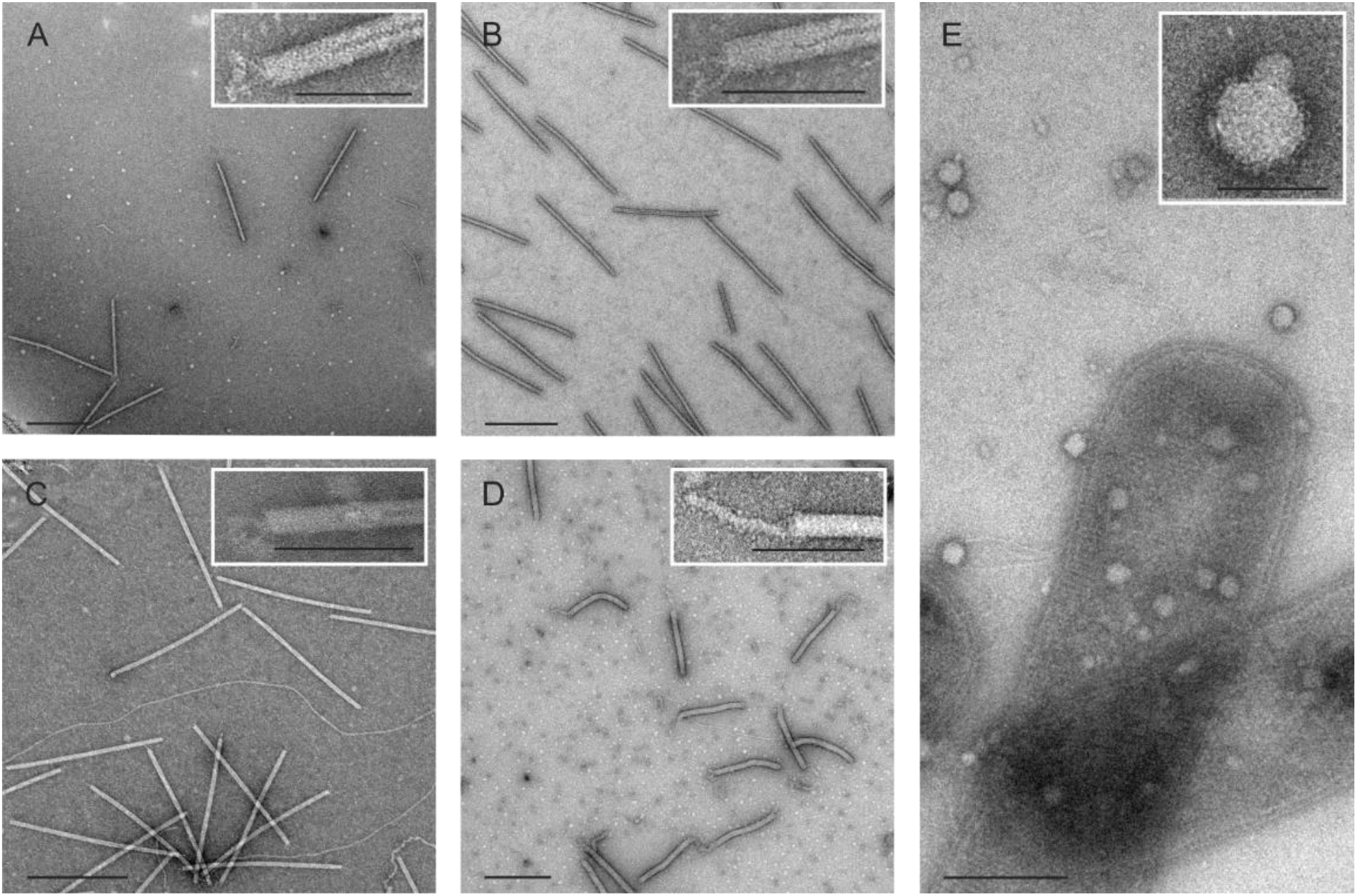
Electron micrographs of the five isolated viruses. A. *Metallosphaera* rod-shaped virus 1. B. *Acidianus* rod-shaped virus 3. C. *Saccharolobus solfataricus* rod-shaped virus 1.D. *Pyrobaculum* filamentous virus 2. E. *Pyrobaculum* spherical virus 2. Samples were negatively stained with 2% (wt/vol) uranyl acetate. Scale bars: 500 nm; in insets: 100 nm.

Negatively stained MRV1, ARV3 and SSRV1 virions are rod-shaped particles measuring 630 ± 20 × 25 ± 2 nm, 670 ± 40 × 23 ± 4 nm and 750 ± 30 × 24 ± 3 nm, respectively (Figure 2A-C). Similar to members of the *Rudiviridae* family, the three viral particles have terminal fibers located at each end of the virion, which have been shown to play a role in host recognition in the case of *Sulfolobus islandicus* rod-shaped virus 2 (SIRV2) [68]. Interestingly, ARV3 virions appear to be more flexible than MRV1 and SSRV1 particles, resembling *Acidianus* rod-shaped virus 1 (ARV1) [69].

A different approach was chosen to identify the hosts of the viral particles detected in the *Pyrobaculum* medium. Exponentially growing liquid cultures of *Pyrobaculum* strains were mixed with concentrated VLPs and incubated for 15 days at 90°C (see Materials and Methods). The replication of the particles was monitored by TEM. *P. arsenaticum* 2GA propagated filamentous and spherical particles, named *Pyrobaculum* filamentous virus 2 (PFV2; Figure 2D) and *Pyrobaculum* spherical virus 2 (PSV2; Figure 2E), respectively. Because *P. arsenaticum* 2GA could not be grown as a lawn on solid medium, dilutions of infected cells were made in order to establish cultures infected with just one type of viral particles.

Negatively stained virions of PFV2 are filamentous and flexible particles of about 450 ± 20 × 34 ± 4 nm in size with terminal filaments of up to 130 nm in length attached to one or both ends of the virions (Figure 2D), similar to what has been reported for PFV1 [45]. PSV2 virions are spherical particles of around 90 ± 20 nm of diameter, with a variable number of bulging protrusions on their surface (Figure 2E), resembling *Pyrobaculum* spherical virus (PSV) particles [70].

Unfortunately, the hosts for other VLPs shown in Figure 1 could not be isolated, either due to unfavorable growth conditions in the laboratory or due to virus-mediated extinction of the corresponding host cell populations.

### Host Range

The host ranges of MRV1, ARV3 and SSRV1 were investigated using strains of the genera *Acidianus, Sulfolobus and Saccharolobus* (Table 1). Only ARV3 could be propagated on *A. hospitalis*, whereas none of the other tested strains served as a host for MRV1 and SSRV1, even though *S. solfataricus* P1 and P2 were also isolated from the solfataric field in Pozzuoli [53]. The host ranges of PFV2 and PSV2 were studied using strains of the genus *Pyrobaculum* (Table 1). None of the tested strains supported propagation of PSV2. By contrast, apparent increase in the number of PFV2 particles was observed by TEM in the presence of *P. oguniense* TE7, albeit not to the same extent as with *P. arsenaticum* 2GA, indicating that *P. oguniense* TE7 is not optimal for the virus propagation.

**Table 1.** Host range of the archaeal viruses isolated in this study.

| Host strain | Newly-isolated archaeal virus |  |  |  |  |
| --- | --- | --- | --- | --- | --- |
|  | MRV1 | ARV3 | SSRV1 | PSV2 | PFV2 |
| *Metallosphaera sedula* POZ202 | H | - | - | - | - |
| *Acidianus brierleyi* POZ9 | - | H | - | - | - |
| *Acidianus convivator* | - | - | - | - | - |
| *Acidianus hospitalis* | - | + | - | - | - |
| *Saccharolobus solfataricus* P1 | - | - | - | - | - |
| *Saccharolobus solfataricus* P2 | - | - | - | - | - |
| *Saccharolobus solfataricus* POZ149 | - | - | H | - | - |
| *Sulfolobus islandicus* LAL14/1 | - | - | - | - | - |
| *Sulfolobus islandicus* REN2H1 | - | - | - | - | - |
| *Sulfolobus islandicus* HVE10/4 | - | - | - | - | - |
| *Sulfolobus islandicus* REY15A | - | - | - | - | - |
| *Sulfolobus islandicus* ΔC1C2 | - | - | - | - | - |
| *Sulfolobus acidocaldarius* DSM639 | - | - | - | - | - |
| *Pyrobaculum arsenaticum* PZ6 (DSM 13514) | - | - | - | - | - |
| *Pyrobaculum arsenaticum* 2GA | - | - | - | H | H |
| *Pyrobaculum calidifontis* VA1 (DSM 21063) | - | - | - | - | - |
| *Pyrobaculum oguniense* TE7 (DSM 13380) | - | - | - | - | + |

H, isolation host; +, supports virus replication.

**Table 2.** Genomic features of the hyperthermophilic archaeal viruses isolated in this study.

| Virus | Host | Size (bp) | TIR | Topology | GC% | ORFs | Accession # |
| --- | --- | --- | --- | --- | --- | --- | --- |
| MRV1 | *Metallosphaera sedula* POZ202 | 20269 | + | linear | 34.12 | 27 | MN876843 |
| ARV3 | *Acidianus brierleyi* POZ9 | 23666 | - | linear | 32.02 | 33 | MN876842 |
| SSRV1 | *Saccharolobus solfataricus* POZ149 | 26097 | - | linear | 32.31 | 37 | MN876841 |
| PSV2 | *Pyrobaculum arsenaticum* 2GA | 18212 | + | linear | 45.01 | 32 | MN876845 |
| PFV2 | *Pyrobaculum arsenaticum* 2GA | 17602 | + | linear | 45.32 | 39 | MN876844 |

TIR, terminal inverted repeats; ORFs, open reading frames.

### Genome organization

Genomes of the five viruses were isolated from the purified virions and treated with DNase I, type II restriction endonucleases (REases) and RNase A. None of the viral genomes were sensitive to RNase A, but could be digested by DNase I and REases, indicating that all viral genomes consist of dsDNA molecules. The genomes were sequenced on Illumina MiSeq platform, with the assembled contigs corresponding to complete or near-complete virus genomes. The general properties of the virus genomes are summarized in Table 2.

#### New species in the Globuloviridae family

The linear dsDNA genome of PSV2 is 18,212 bp in length and has a GC content of 45%, which is similar to that of other *Pyrobaculum*-infecting viruses (45-48%) [45,70]. The coding region of the PSV2 genome is flanked by perfect 55 bp-long terminal inverted repeats (TIR), confirming the linear topology and (near) completeness of the genome. It contains 32 predicted open reading frames (ORFs), all located on the same strand. Thirteen PSV ORFs contain at least one predicted membrane-spanning region. Notably, two of them (ORFs 3 and 9) have nine predicted transmembrane domains.

Globuloviruses stand out as some of the most mysterious among archaeal viruses, with 98% of their proteins showing no similarity to sequences in public databases [25]. In part, this is due to the fact that the family is represented by only two members, PSV and TTSV1, infecting *Pyrobaculum* [70] and *Thermoproteus tenax* [71], respectively, which precludes generation of sensitive sequence profiles for remote homology searches. Previous structural genomics initiatives were also not very illuminating [72]. As a result, the vast majority of the globulovirus proteins lack functional annotation. Homologs of PSV2 proteins were identified exclusively in members of the *Globuloviridae* (Figure 3), corroborating the initial affiliation of PSV2 into the family *Globuloviridae* based on the morphological features of the virion. Nineteen PSV2 ORFs, including those encoding the three major structural proteins (VP1-VP3), have closest homologs in PSV [70] with amino acid sequence identities ranging between 28% and 65% (Figure 3); 13 of these ORFs are also shared with TTSV1. The remaining 13 PSV2 ORFs yielded no significant matches to sequences in public databases. The three genomes display no appreciable similarity at the nucleotide sequence level, indicating considerable sequence diversity within the *Globuloviridae* family. Notably, among five PSV proteins for which high-resolution structures are available [72], only one protein with a unique fold, PSV gp11 (ORF239), is conserved in PSV2.

**Figure 3.**
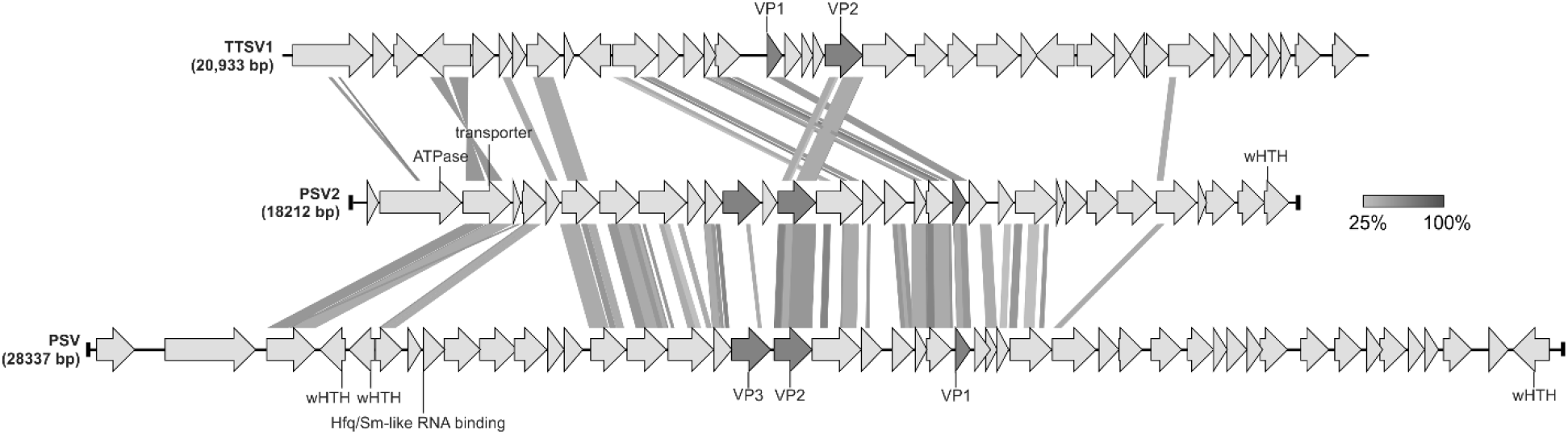
Genome alignment of the three members of the *Globuloviridae* family. The open reading frames (ORFs) are represented by arrows that indicate the direction of transcription. The terminal inverted repeats (TIRs) are denoted by black bars at the ends of the genomes. Genes encoding the major structural proteins are shown in dark grey. The functional annotations of the predicted ORFs are depicted above/below the corresponding ORF. Homologous genes are connected by shading in grayscale based on the level of amino acid sequence identity. VP, virion protein; wHTH, winged helix-turn-helix domain.

Sensitive profile-profile comparisons using HHpred allowed functional annotation of only four PSV2 proteins. The PSV2 ORF2 encodes a protein with an AAA+ ATPase domain, which is most closely related to those found in ClpB-like chaperones and heat shock proteins (HHpred probability of 99.6%; Supplementary Information, Table S2). ORF3 encodes one of the two proteins with nine putative transmembrane domains and is predicted to function as a membrane transporter, most closely matching bacterial and archaeal cation exchangers (HHpred probability of 95.5%). Interestingly, the product of ORF4 is predicted to be a circadian clock protein KaiB, albeit with a lower probability (HHpred probability of 93%). Finally, ORF32 shares homology with a putative transcriptional regulator with the winged helix-turn-helix domain (wHTH) of *Saccharolobus solfataricus* (HHpred probability of 94.4%). In addition, ORF11 encodes a functionally uncharacterized DUF1286-family protein conserved in archaea and several *Saccharolobus*-infecting viruses (Supplementary Information, Table S2).

#### New member in the Tristromaviridae family

The linear genome of the isolated PFV2 carries 17,602 bp and contains 39 ORFs, all except one located on the same strand. The coding region is flanked by 59 bp-long TIRs. The GC content (45.3%) of the genome is similar to that of PSV2 and other *Pyrobaculum-*infecting viruses [45,70], but is considerably lower than in *Pyrobaculum arsenaticum* PZ6 (58.3%), and *Pyrobaculum oguniense*TE7 (55.1%). Eleven of the PFV2 ORFs were predicted to encode proteins with one or more membrane-spanning domains (Supplementary Information, Table S2).

The PFV2 genome is 98.9% identical over 70% of its length to that of PFV1, the type species of the *Tristromaviridae* family [45]. PFV1 and PFV2 were isolated ~3 years apart, from the same solfataric field in Pozzuoli [45], suggesting that the population of tristromaviruses is relatively stable over time. Accordingly, 36 of the 39 PFV2 ORFs are nearly identical (average amino acid identity of 96.5%) to those of PFV1. Two events account for the differences between PFV1 and PFV2: (i) a deletion spanning most of the PFV1 gene 27 (including codons 72-495) as well as the downstream genes 28-30, and (ii) insertion of a four-gene block between PFV1 genes 36 and 37 in PFV2 (Figure 4). PFV1 gene 27 encodes a minor virion protein, whereas genes 29 and 30 encode putative small, lectin-like carbohydrate-binding proteins (Rensen et al., 2016). The absence of the corresponding genes in PFV2 genome suggests that they are dispensable for the PFV1/PFV2 infection cycle. Notably, a homolog of the PFV1 glycoside hydrolase gene 28 is reinserted into PFV2 genome as part of the four-gene block (PFV2 ORF34). However, the two genes do not appear to be orthologous in PFV1 and PFV2 genomes, because they share much lower sequence similarity (50% versus average 96.5% identity) compared to other orthologous genes. By contrast, PFV2 ORF35 has no counterpart in PFV1 but is homologous to the glycosyltransferase gene of *Thermoproteus tenax* virus 1 (TTV1) [73], the only other known member of the *Tristromaviridae* family [74](Figure 4). Thus, comparison of the closely related tristromavirus sequences revealed active genome remodeling in this virus group, involving both deletions and horizontal acquisition of new genes.

**Figure 4.**
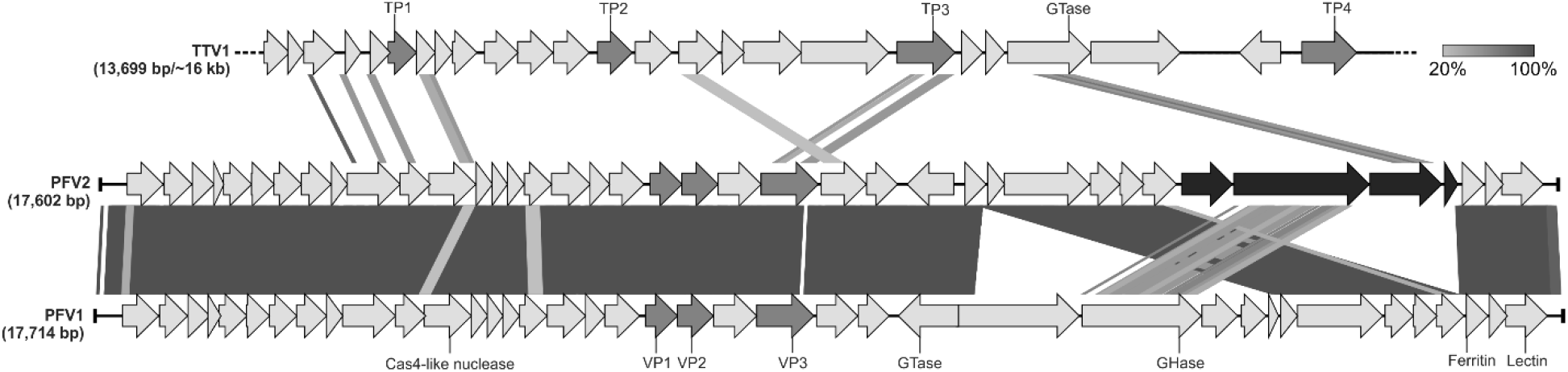
Genome comparison of the three members of the *Tristromaviridae* family. The open reading frames (ORFs) are represented by arrows that indicate the direction of transcription. The terminal inverted repeats (TIRs) are denoted by black bars at the ends of the genomes. Genes encoding the major structural proteins are shown in dark grey, whereas the four-gene block discussed in the text is shown in black. The functional annotations of the predicted ORFs are depicted above/below the corresponding ORFs. Homologous genes are connected by shading in grayscale based on the level of amino acid sequence identity. The dotted line represents the incompleteness of the TTV1 genome. GHase, glycoside hydrolase; GTase, glycosyltransferase; TP/VP, virion protein.

#### New rod-shaped viruses

The linear genomes of the isolated rudiviruses have a *length* ranging from 20,269 to 26,079 bp and a GC content varying between 32.02% and 34.12%. The MRV1 genome contains 80 bp-long TIR, suggesting that the genome is coding-complete (i.e., contains all protein-coding genes). Although no TIRs could be identified for ARV3 and SSRV1, comparison with the genomes of other rudiviruses (Figure 5) suggests that the two genomes are also nearly complete.

**Figure 5.**
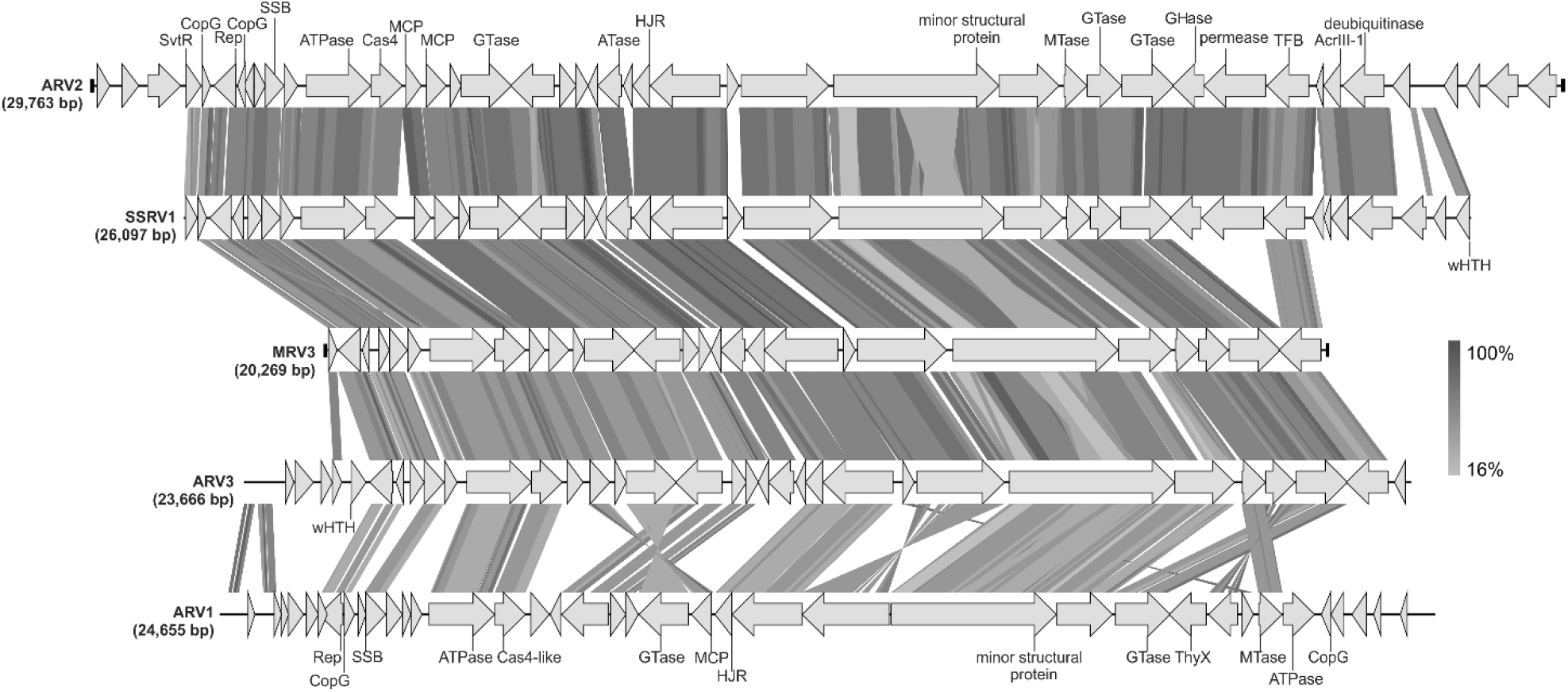
Genome alignment of the Italian rudiviruses. ARV3, MRV1 and SSRV1 are newly isolated members of the *Rudiviridae* family, whereas ARV1 and ARV2 were reported previously [69,75]. The open reading frames (ORFs) are represented by arrows that indicate the direction of transcription. The terminal inverted repeats (TIRs) are denoted by black bars at the ends of the genomes. The functional annotations of the predicted ORFs are depicted above/below the corresponding ORFs. Homologous genes are connected by shading in grayscale based on the amino acid sequence identity. ATase, acetyltransferase; GHase, glycoside hydrolase; GTase, glycosyltransferase; HJR, Holliday junction resolvase; MCP, major capsid protein; MTase, methyltransferase; SSB, ssDNA binding protein; TFB, Transcription factor B; ThyX, thymidylate synthase; wHTH, winged helix-turn-helix domain.

The genomes of MRV1, ARV3 and SSRV1 contain 27, 33 and 37 ORFs, respectively, and display high degree of gene synteny (Figure 5). Comparison of the three genomes showed that they share 26 putative proteins, with amino acid sequence identities ranging between 35.1% and 93.9%. Among the previously reported rudiviruses, the three newly isolated viruses share the highest similarity with ARV2 (ANI of ~78%), which was metagenomically sequenced from samples collected in the same Pozzuoli area [75]; this group of viruses shares 20 genes.

Rudiviruses represent one of the most extensively studied families of archaeal viruses with many of the viral proteins being functionally and structurally characterized [76]. Most of the MRV1, ARV3 and SSRV1 ORFs have orthologs in at least some members of the *Rudiviridae*. Genes shared with other rudiviruses include those for the major and minor capsid proteins, several transcription factors with ribbon-helix-helix motifs, three glycosyltransferases, GCN5-family acetyltransferase, Holliday junction resolvase, SAM-dependent methyltransferase and a gene cassette encoding the ssDNA-binding protein, ssDNA annealing ATPase and Cas4-like ssDNA exonuclease (Figure 5). The conservation of these core proteins in the expanding collection of rudiviruses underscores their critical role in viral reproduction. Conspicuously missing from the MRV1 and ARV3 are homologs of the SIRV2 P98 protein responsible for the formation of pyramidal structures for virion egress and anti-CRISPR (Acr) proteins encoded by diverse crenarchaeal viruses [77,78]. Notably, SSRV1 carries a gene for the recently characterized AcrIII-1, which blocks antiviral response of type III CRISPR-Cas systems by cleaving the cyclic oligoadenylate second messenger [79]. Given that CRISPR-Cas systems are highly prevalent in hyperthermophilic archaea[80], the lack of identifiable anti-CRISPR genes in MRV1 and ARV3 is somewhat unexpected, suggesting that the two viruses encode novel Acr proteins. Similarly, lack of the genes encoding recognizable P98-like pyramid proteins in MRV1 and ARV3 (as well as in ARV1 and ARV2) suggests that these viruses have evolved a different solution for virion release.

Besides the core genes, rudiviruses are known to carry a rich complement of variable genes, which typically occupy the termini of linear genomes and are shared with viruses isolated from the same geographical location [81]. For instance, MRV1, ARV3 and SSRV1 carry several genes, which are exclusive to Italian rudiviruses. These include a divergent glycoside hydrolase, putative metal-dependent deubiquitinase, alpha-helical DNA-binding protein, transcription initiation factor and several short hypothetical proteins, which are likely candidates for Acrs (Supplementary Information, Table S2). Notably, some of these hypothetical proteins are shared with other crenarchaeal viruses (families *Lipothrixviridae* and *Bicaudaviridae*) isolated from the same location.

### A biogeographic pattern in the *Rudiviridae* family

To gain insight into the global architecture of the rudivirus populations and the factors that govern it, we performed phylogenomic analysis of all available rudivirus genomes using the Genome-BLAST Distance Phylogeny method implemented in VICTOR [66]. Our results unequivocally show that the 19 sequenced rudiviruses fall into six clades corresponding to the geographical origins of the virus isolation (Figure 6), suggesting local adaptation of the corresponding viruses. Thus, on the global scale, horizontal spread of rudivirus virions between geographically remote continental hydrothermal systems appears to be restricted.

**Figure 6.**
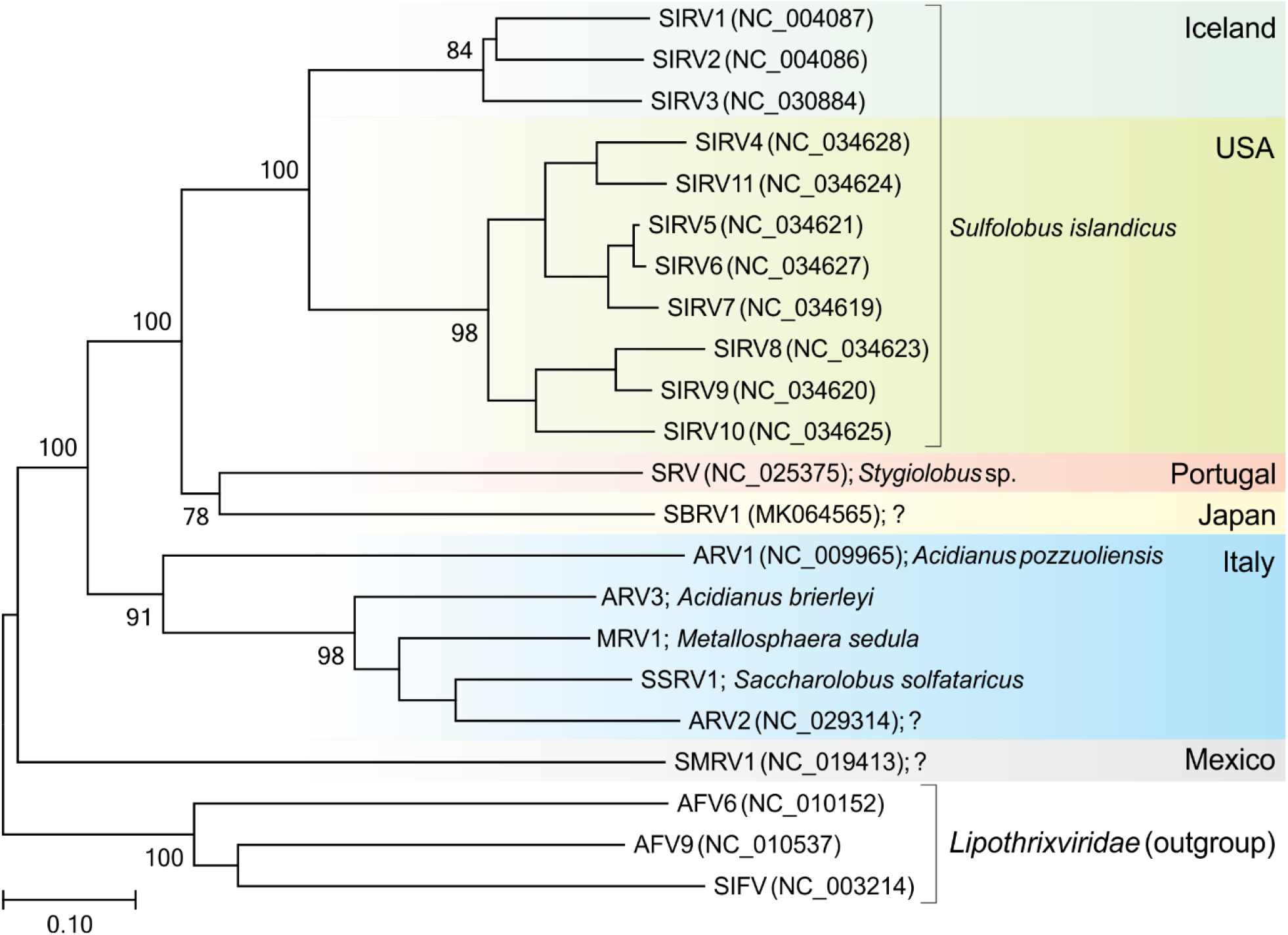
Inferred phylogenomic tree of all known members of the *Rudiviridae* family based on whole genome VICTOR [66] analysis at the amino acid level. The tree is rooted with lipothrixviruses, and the branch length is scaled in terms of the Genome BLAST Distance Phylogeny (GBDP) distance formula D6. Only branch support values >70% are shown. For each genome, the abbreviated virus name, GenBank accession number and host organism (when known) are indicated. Question marks denote that the host is not known. The tree is divided into colored blocks according to the geographical origin of the compared viruses.

Remarkably, despite forming a monophyletic group, all rudiviruses originating from Italy infect relatively distant hosts, belonging to three different genera of the order Sulfolobales. Such pattern was highly unexpected, because closely related hyperthermophilic archaeal viruses typically infect hosts from the same genus. Thus, we hypothesized that such pattern of host specificities might signify host switching events in the history of the Italian rudivirus assemblage. CRISPR arrays, which contain spacer sequences derived from mobile genetic elements, keep memory of past infections and are commonly used as indicators for matching the uncultivated viruses to their potential hosts [82–84]. Thus, to investigate which hosts were exposed to Italian rudiviruses, we searched the CRISPRdb database [85] for the presence of spacers matching the corresponding viral genomes. Spacer matches were found for all five virus genomes, albeit with different levels of identity. Matches with 100% identity were obtained for ARV2, ARV3, SSRV1 and MRV1 (Supplementary Information, Table S3). Spacers matching ARV2 and ARV3 were found in *Saccharolobus solfataricus* P1, whereas ARV2 was also targeted by a CRISPR spacer (100% identity) from *Metallosphaera sedula* DSM 5348. Unexpectedly, MRV1, which infects *Metallosphaera* species, was matched by spacers from different strains of *S. solfataricus*. Conversely, SSRV1 infecting *S. solfataricus* was targeted by multiple spacers from CRISPR arrays of *M. sedula* DSM 5348 (Supplementary Information, Table S3). These results are consistent with the possibility that in the recent history of Italian rudiviruses, host switching, even across the genus boundary, has been relatively common.

### Revised classification of rudiviruses

Of the 19 rudiviruses for which (near) complete genome sequences are available, only three (SIRV1, SIRV2 and ARV1) are officially classified. All three viruses are included in the same genus, *Rudivirus*. Here, we propose a taxonomic framework for classification of all cultivated and uncultured rudiviruses for which genome sequences are available. As mentioned above, phylogenomic analysis revealed six different clades coinciding with the origin of virus isolation (Figure 6), highlighting considerable diversity of the natural rudivirus population, which has stratified into several assemblages, warranting their classification into distinct, genus-level taxonomic units. To this end, we compared the genome sequences using the Gegenees tool [67], which fragments the genomes and calculates symmetrical identity scores for each pairwise comparison based on BLASTn hits and a genome length. The analysis revealed seven clusters of related genomes, which were generally consistent with those obtained in the phylogenomic analysis (Table S4). Notably, due to considerable sequence divergence, ARV1 falls into a separate cluster from other Italian rudiviruses. Consistently, in the phylogenomic tree, ARV1 forms a sister group to other Italian rudiviruses. Furthermore, among the five Italian rudiviruses, the genome of ARV1 is most divergent, displaying multiple gene and genomic segment inversions compared to the other viruses (Figure 5). Viruses from the seven clades also differ considerably in terms of the variable gene contents. For instance, viruses SIRV1 and SIRV2 isolated in Iceland share 11 genes that are absent from the USA SIRVs [81]. Thus, to acknowledge the differences between the known rudiviruses, we propose to classify them into seven genera: “*Icerudivirus*” (former *Rudivirus*,to include SIRV1-SIRV3), “*Mexirudivirus*” (SMRV1), “A*zorudivirus*” (SRV), “*Itarudivirus*” (ARV1), “*Hoswirudivirus*” (ARV2, ARV3, MRV1, SSRV1; *hoswi-*, for <u>ho</u>st <u>swi</u>tching), “*Japarudivirus*” (SBRV1) and “*Usarudivirus*” (SIRV4-SIRV11).

## DISCUSSION

Virus discovery in the “age of metagenomics” is increasingly performed by culture-independent methods [86], whereby viral genomes are sequenced directly from the environment. This is a powerful approach which has already yielded thousands of viral genomes, providing unprecedented insights into virus diversity, environmental distribution and evolution [82,87–90]. The limitation of viral metagenomics, however, is that the host species for the sequenced viruses typically remain unknown, even if occasionally predicted. Furthermore, truly novel viruses, which lack similarity to known reference virus genomes, often escape identification and remain in the so-called “viral dark matter” [91–94]. Finally, although many features of viruses can be deduced from the genome sequence alone, different molecular aspects of virus-host interactions remain obscure. Thus, to get a full(er) picture of virus biology and to further expand the knowledge on virus diversity, classical approaches of virus isolation and characterization remain highly valuable. Here, we reported the results of our exploration of the diversity of hyperthermophilic archaeal viruses at the solfataric field in Pisciarelli, Pozzuoli.

Previous sampling of archaeal viruses in the thermal springs in Pisciarelli led to the isolation of viruses from the families *Ampullaviridae*, *Bicaudaviridae*, *Lipothrixviridae*, *Rudiviridae* and *Tristromaviridae* [31,45,69,75,95], whereas those of the families *Fuselloviridae*, *Globuloviridae* and *Guttaviridae*, which we observed in the initial enrichment cultures, have not been previously reported from the sampled Pisciarelli sites. Nevertheless, similar virion morphologies have been observed in high-temperature continental hydrothermal systems from other geographical locations across the globe [29,32,70,96,97], pointing to global distribution of most groups of hyperthermophilic archaeal viruses. However, how these virus communities are structured and whether geographically remote hydrothermal ecosystems undergo virus immigration is not fully understood.

A previous study has revealed a biogeographic pattern among *Sulfolobus islandicus*-infecting rudiviruses isolated from hot springs in Iceland and United States [81]. That is, viruses from the same geographical location were more closely related to each other than they were to viruses from other locations and the larger the distance the more divergent the virus genomes were. Similar observation has been made for the *Sulfolobus*-infecting spindle-shaped viruses of the family *Fuselloviridae* [97–99]. By contrast, a study focusing on three relatively closely spaced hot springs in Yellowstone National Park concluded that horizontal virus movement, rather than mutation, is the dominant factor controlling the viral community structure [100].

Phylogenomic and comparative genomic analysis of the rudiviruses reported herein and those sequenced previously revealed a strong biogeographic pattern. All rudiviruses formed clades corresponding to the geographical origins of their isolation. This observation strongly suggests that diversification and evolution of rudiviruses is influenced by special confinement within discrete high-temperature continental hydrothermal systems, with little horizontal migration of viral particles over large distances. Consistently, analyses of the CRISPR spacers carried by hyperthermophilic archaea predominantly target local viruses, further indicating geographically defined co-evolution of viruses and their hosts [97,101]. This is in stark contrast with the global architecture of virus communities associated with hyperhalophilic archaea, where genomic similarity between viruses does not correspond to geographical distance [102,103]. Indeed, it has been suggested that hypersaline environments worldwide function like a single habitat, with extensive exchange of microbes and their viruses between remote hypersaline sources [102]. It has been shown that small salt crystals harboring halophilic archaea are carried on bird feathers, suggesting that bird migration is a driving force of haloarchaeal spread across the globe [104]. Obviously, similar route of dissemination is hardly imaginable for microbial communities thriving in acidic high-temperature hydrothermal settings, offering an explanation for the different patterns observed for hyperthermophilic and hyperhalophilic archaeal viruses. Notably, however, it has been suggested that reversible silicification of virus particles, which is conceivable in hot spring environments, might promote long-distance host-independent virus dispersal [105].

We also show, for the first time, that relatively closely related rudiviruses infect phylogenetically distant hosts, belonging to three different genera. This observation is inconsistent with stable coevolution of rudiviruses with their hosts, but rather points to host switch events in the recent history of Italian rudiviruses. At least in the case of rudivirises, relatively few genetic changes appear to be necessary for gaining the ability to infect a new host. In tailed bacteriophages, host range switches are typically associated with mutations in genes encoding the tail fiber proteins responsible for host recognition [106]. Several molecular mechanisms underlying mutability of the tail fiber genes have been described, including genetic drift, diversity generating retroelements and phase variation cassettes. The latter two systems have been demonstrated to function also in viruses infecting anaerobic methane-oxidizing (ANME) and hyperhalophilic archaea, respectively [107,108]. Both mechanisms depend on specific enzymes, namely, reverse-transcriptase and invertase. However, neither of the two systems is present in rudiviruses, suggesting that genetic drift is the most likely mechanism responsible for generating diversity in the host recognition module of rudiviruses.

Analysis of CRISPR spacers from hyperthermophilic archaea confirmed that viruses closely related to those isolated in this study were infecting highly different hosts in the recent past. Indeed, viruses infecting *Saccharolobus* and *Metallosphaera* species were targeted by spacers from CRISPR arrays of *Metallosphaera* and *Saccharolobus*, respectively. This observation further reinforces a relatively recent host switch event. Notably, this phenomenon appears to be also applicable to rudivirus from other geographical locations. In particular, SBRV1, a rudivirus from a Japanese hot spring [109], was found to be targeted by 521 unique CRISPR spacers associated with all four principal CRISPR repeat sequences present in Sulfolobales [101], suggesting either very broad host range or frequent host switches. Furthermore, we have recently shown that CRISPR targeting is an important factor driving the genome evolution of hyperthermophilic archaeal viruses [101]. Given the presence of multiple CRISPR spacers matching rudivirus genomes, we hypothesize that necessity to switch hosts might be, to a large extent, driven by CRISPR targeting. Notably, matching of CRISPR spacers to protospacers in viral genomes is one of the most widely used approaches of host identification for viruses discovered by metagenomics, with the estimated host genus prediction accuracy of 70–90% [83,84,89]. Our results suggest that in the case of rudiviruses, spacer matching might not provide accurate predictions beyond the rank of family (i.e., Sulfolobaceae).

Collectively, we show that terrestrial hydrothermal systems harbor a highly diverse virome represented by multiple families with unique virus morphologies not described in other environments. Genomes of the newly isolated viruses, especially those infecting *Pyrobaculum* species, remain a rich source of unknown genes, which could be involved in novel mechanisms of virus-host interactions. Furthermore, our results suggest that global rudivirus communities display biogeographic pattern and diversify into distinct lineages confined to discrete geographical locations. This diversification appears to involve relatively frequent host switching, potentially evoked by host CRISPR-Cas immunity systems. Future studies should focus on characterization of molecular mechanisms allowing rudiviruses to avoid CRISPR targeting by infecting new hosts.

## ACKNOWLEDGEMENTS

This work was supported by l’Agence Nationale de la Recherche (France) project ENVIRA (to M.K.) and the European Union’s Horizon 2020 research and innovation program under grant agreement 685778, project VIRUS X (to D.P.). D.P.B. is part of the Pasteur – Paris University (PPU) International PhD Program. This project has received funding from the European Union’s Horizon 2020 research and innovation programme under the Marie Sklodowska-Curie grant agreement No 665807. We are also grateful to the Ultrastructural BioImaging (UTechS UBI) unit of Institut Pasteur for access to electron microscopes and Marc Monot from the Biomics Platform of Institut Pasteur for helpful discussions on genome assembly.

